# TractoFlow-ABS (Atlas-Based Segmentation)

**DOI:** 10.1101/2020.08.03.197384

**Authors:** Guillaume Theaud, Jean-Christophe Houde, Arnaud Boré, François Rheault, Felix Morency, Maxime Descoteaux

## Abstract

In Diffusion MRI (dMRI), pathological brains are a challenge for tractography processing, where most pipelines are not are not robust to white matter lesions. Intensity of white matter lesions on T1 images can have similar contrasts to gray matter tissue, which leads to misclassifications or “holes” in the white matter mask. These holes produce premature stop for tracking algorithms. To handle these issues, we developed *TractoFlow-ABS* (<u>A</u>tlas-<u>B</u>ased <u>S</u>egmentation). *TractoFlow-ABS* uses the *Freesurfer* atlas to compute tissue masks instead of *FSL fast* in standard *TractoFlow*. *TractoFlow-ABS* is therefore a derived version of *TractoFlow* that is robust to white matter anomalies such as hyperintensities and lesions.

## 1. Introduction

White matter can be affected by multiple types of lesions due to aging and multiple sclerosis, amongst other diseases. In the field of diffusion MRI, these lesions can affect DWI processing and more precisely the tractography step [Edde et al., 2019; Theaud et al., 2017]. In order to do the best processing as possible, a version of *TractoFlow* [Theaud et al., 2020] was developed for brains where the usual T1 segmentation step fails. Compared to *TractoFlow*, *TractoFlow-ABS* uses Freesurfer to compute tissue masks instead of *fast* from FSL. Lesion instensities on T1 images are similar to gray matter tissue, which leads to misclassifications from *fast*. Using Freesurfer allows to have a better tissue classification, which then results in better tracking masks and a better tractogram. Otherwise, the tractograpy process stops prematureraly due the holes and the misclassified gray matter voxels in the deep white matter.

## 2. Methods

*TractoFlow-ABS* is based on *TractoFlow* [Theaud et al., 2020].

### Pipeline inputs

*TractoFlow-ABS* requires a Freesurfer [Fischl, 2012] output computed from the original T1 in native space. Then, the pipeline takes as input: the diffusion weighted images, T1 weighted image, b-values and b-vectors, wmparc and aparc+aseg from Freesurfer. As in *TractoFlow*, if available, the pipeline accepts a reversed phase encoded b=0 image. These input files are illustrated in Figure 1.A.

**Figure 1:**
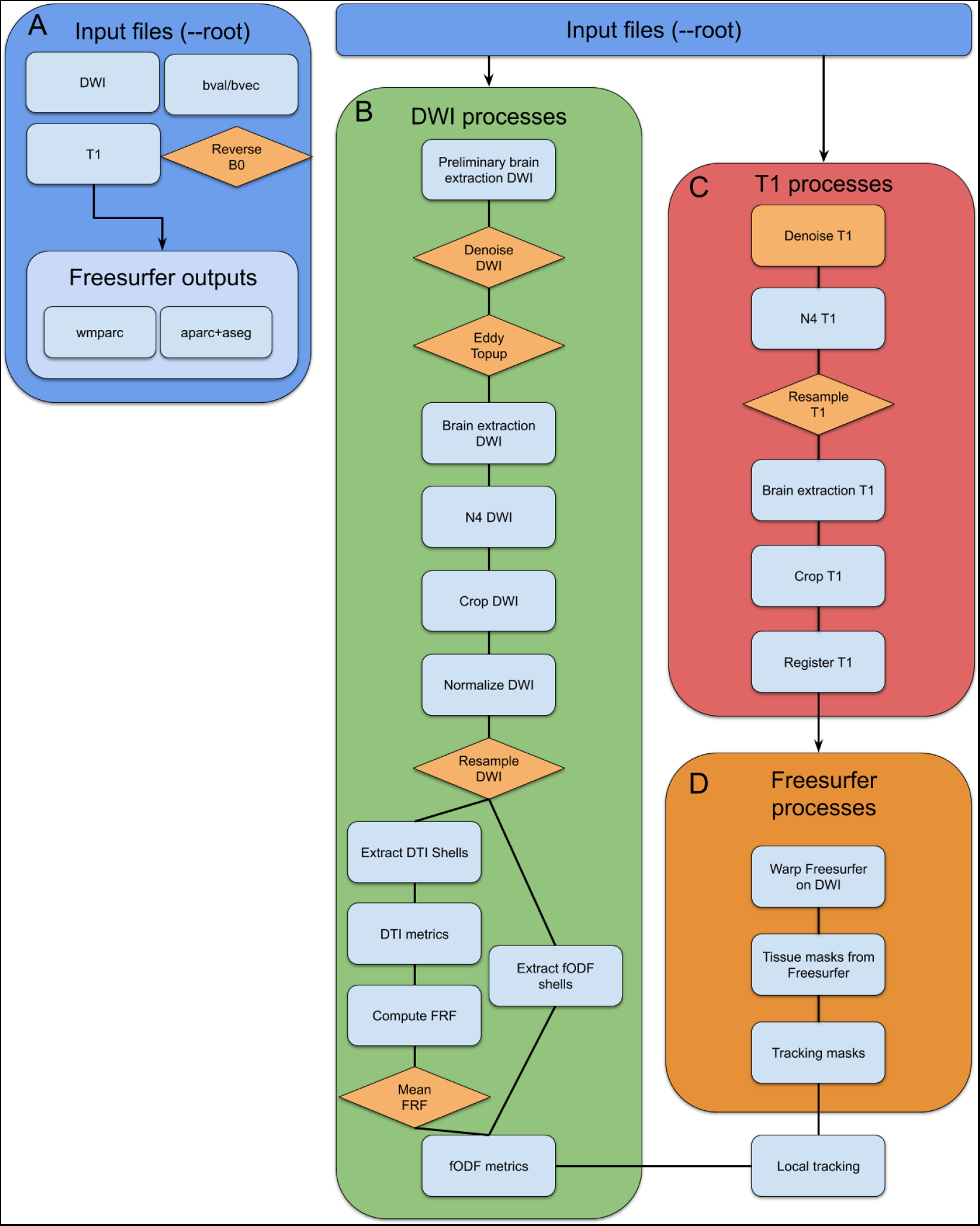
The graph of processes of the *TractoFlow-ABS* pipeline. In A (blue), the input files required to run the pipeline. In B (green), the DWI processes that take the DWI, the b-values/b-vectors files and the reversed phase encoded b=0 image. In C (red), the T1 processes that take the T1-weighted image as input. In D, the Freesurfer processes that take wmparc and aparc+aseg. In orange, all processes or images that are optional.

### Diffusion-weighted images (DWI) tasks

The DWI tasks are the same as presented in the original *TractoFlow* paper [Theaud et al., 2020]. The steps are illustrated in Figure 1.B.

### T1-weighted image tasks

T1-weighted image tasks are similar to *TractoFlow* except for the tissue maps and the tracking maps steps, as seen in 1.C. As said in the introduction, with aging or multiple sclerosis, white matter lesions appear and can impact results of tractography algorithms by creating holes in the tracking mask. In order to fill in the holes due to white matter lesions, some Freesurfer tasks are added to *TractoFlow-ABS*. It is important to point out that one could decide to use this strategy for healthy brains.

### Freesurfer tasks

Freesurfer tasks consist of 3 steps (1.D). First, Freesurfer wmparc and aparc+aseg output files are warped on the DWI using the nonlinear transformation from the T1-to-Diffusion registration step, with nearestneighbor interpolation. Then, WM (white matter), GM (gray matter), CSF (cerebral spinal fluid) and nucleii masks are extracted from the Freesurfer images combining the appropriate labels (see Figure 2). Finally, tracking and seeding masks are created.

**Figure 2:**
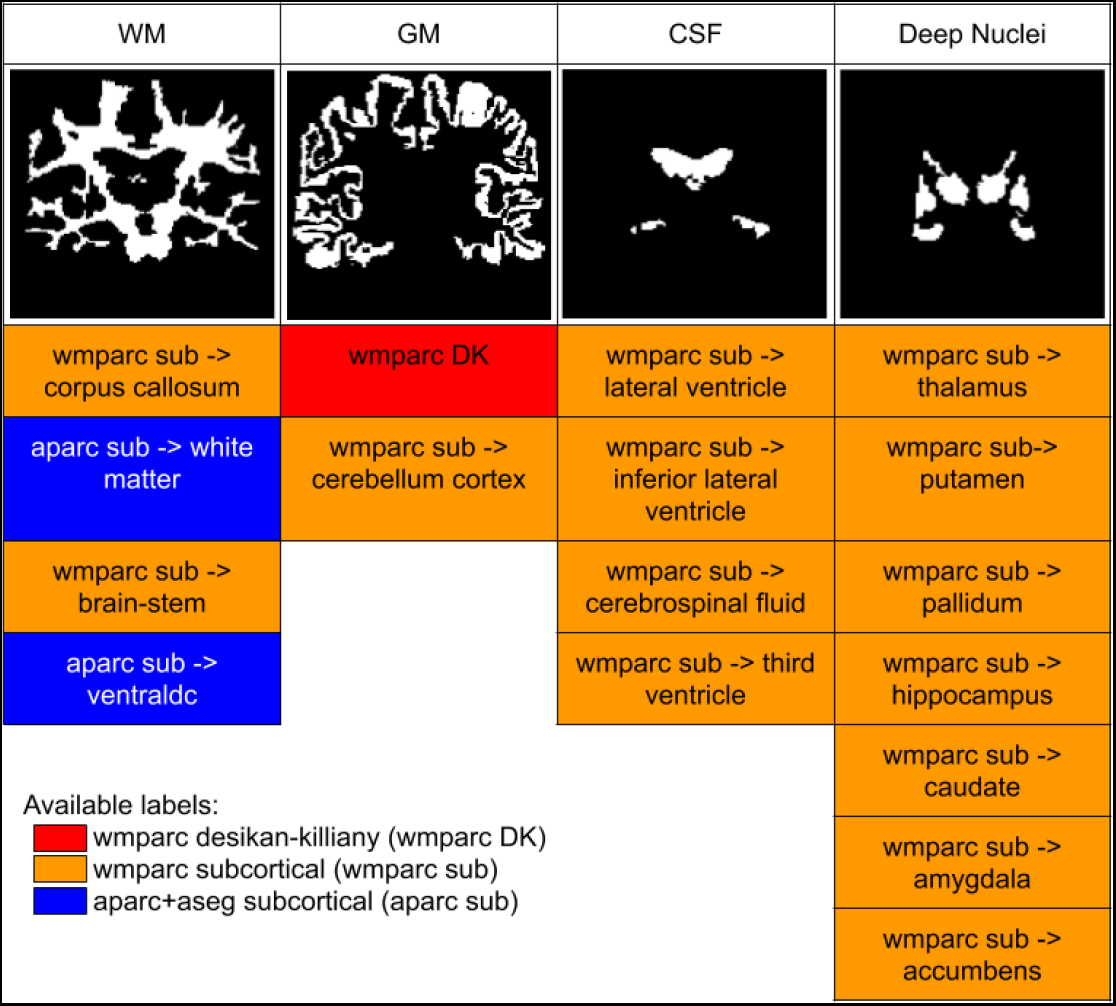
Details of Freesurfer label combining.

### Tracking task

For *TractoFlow-ABS*, the tracking algorithm used is the local tracking [Descoteaux et al., 2009], as opposed to the particle filter tractography (PFT) algorithm used in *TractoFlow*, which requires probabilistic tissue maps. As in *TractoFlow*, step size, number of seeds per voxel, deterministic or probabilistic algorithms can be selected by the user with command-line options.

## 3. Results

In this part, *TractoFlow* and *TractoFlow-ABS* are compared on a simple illustrative example of tissue maps and tractography output of the two pipelines using a subject from the ADNI database.

### Tissue maps comparisons

Tissue maps segmented from *TractoFlow* are impacted by the WM lesions. Lesions are classified as GM instead of WM, as seen in Figure 3. This misclassification has a big impact on tracking masks, which are derived from theses tissue maps. Impact on tractography will be seen in the next section.

**Figure 3:**
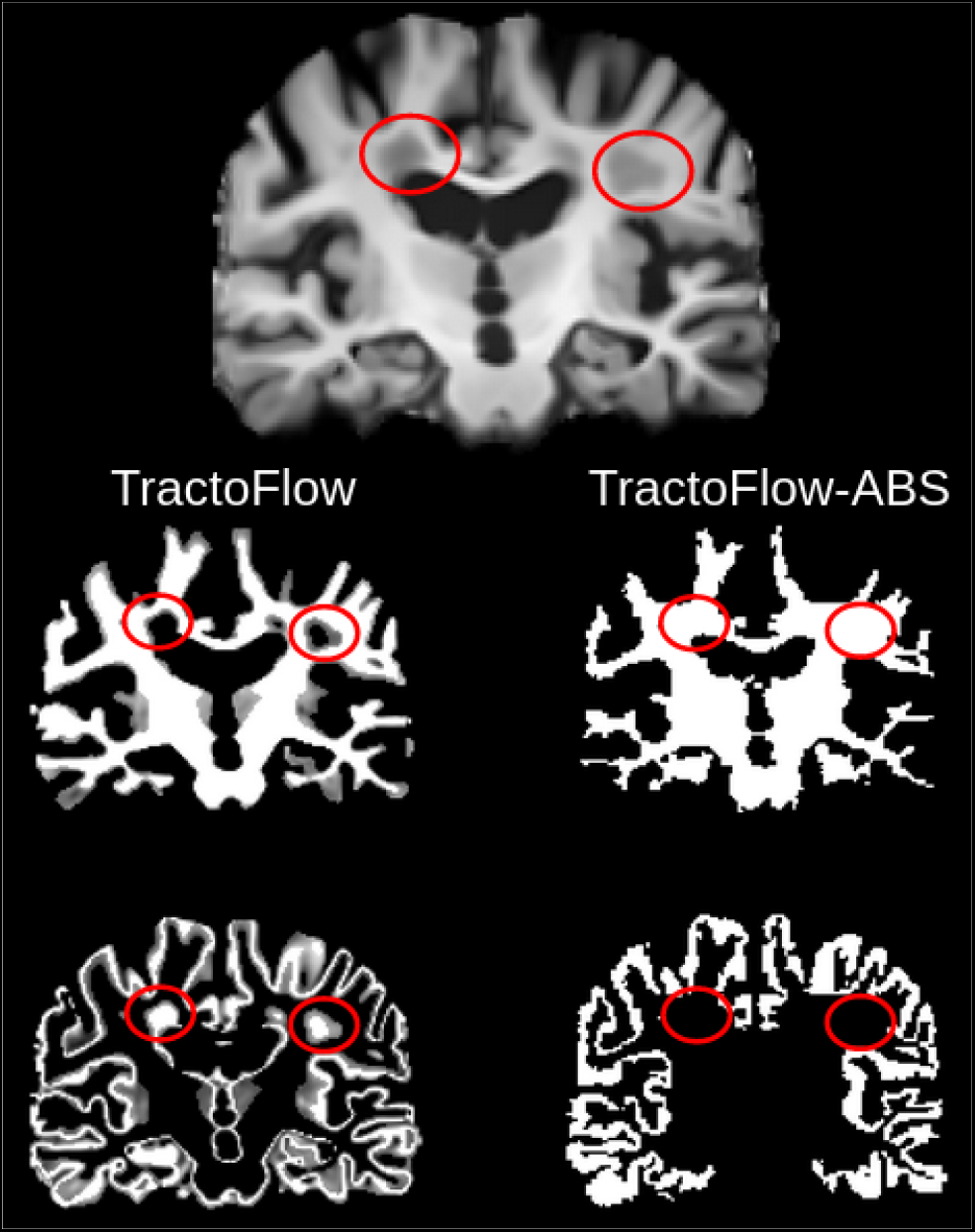
In red circles, lesions locations. In the left column, tissue maps results from *TractoFlow*. In the right column, tissue maps from *TractoFlow-ABS*

**Figure 4:**
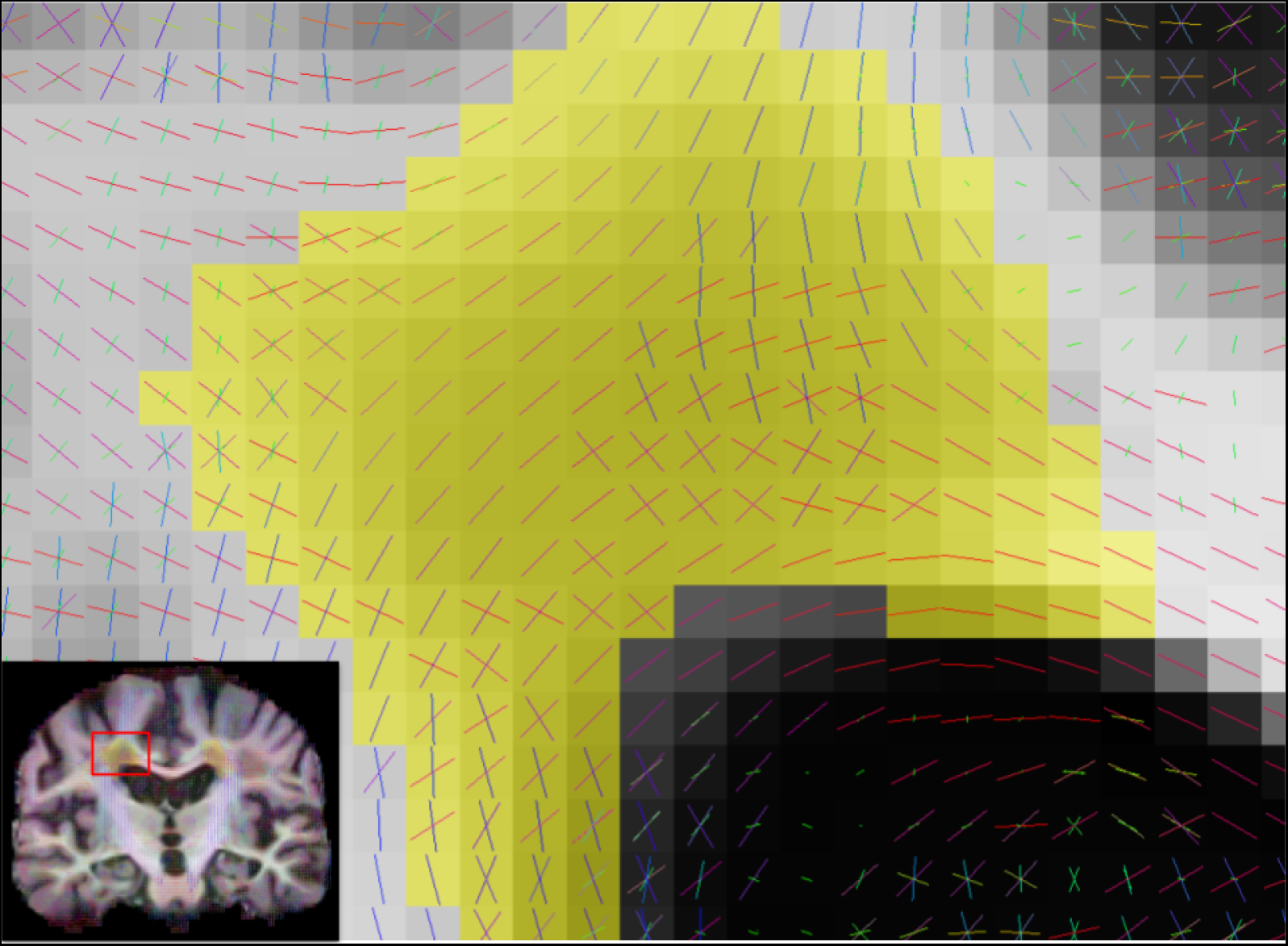
In yellow, a white matter lesion. Local orientations in the lesions are coherent. Local tractography can thus reconstruct pathway through this lesion.

With *TractoFlow-ABS*, this misclassification is solved by the freesurfer atlas, and the WM lesions are classified as WM and not GM. This improvement will permit the tractography algorithm to explore this region and reconstruct pathways through these lesions, if coherent local orientations are present (as seen in Figure 5).

**Figure 5:**
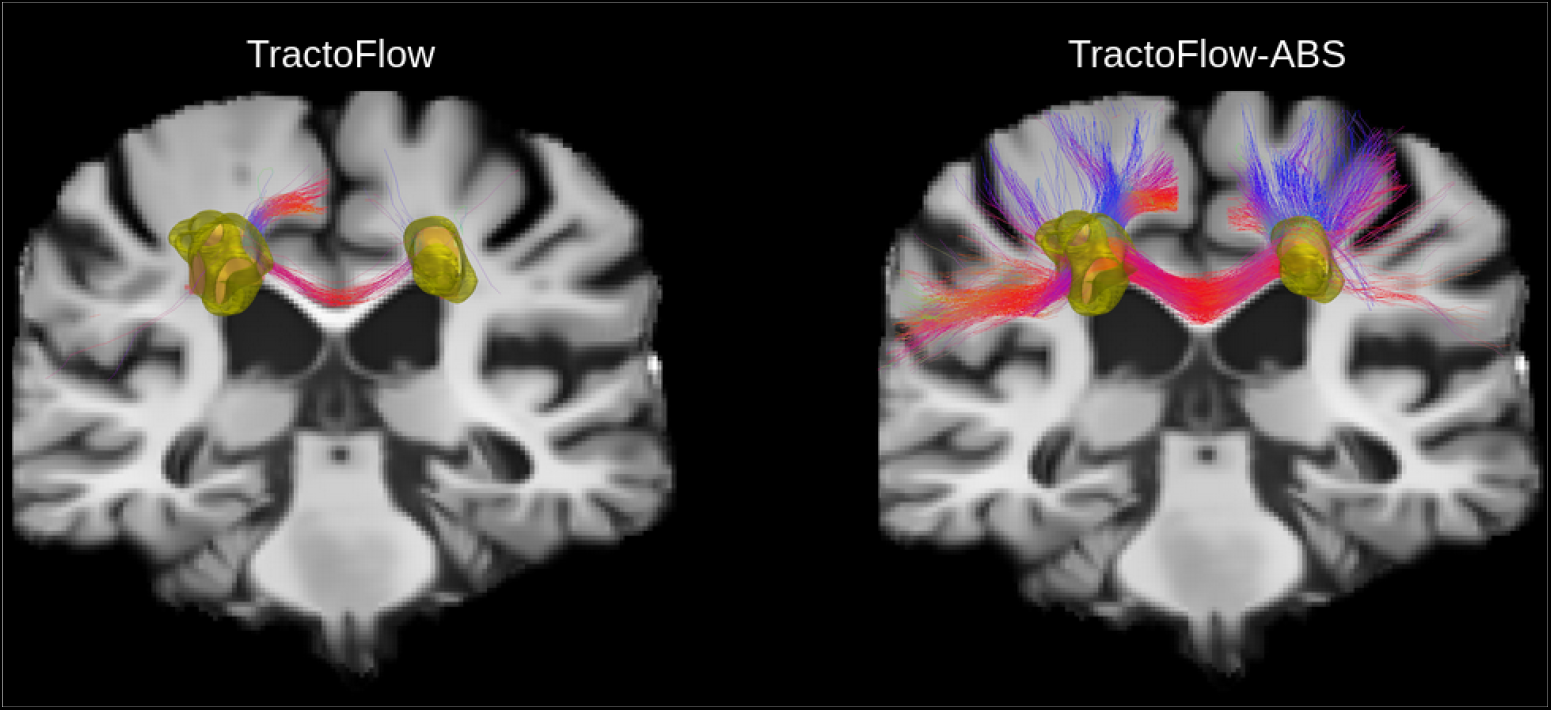
The corpus callosum extracted from *TractoFlow* and *TractoFlow-ABS* tractograms showed in a coronal slice view. In yellow, WM lesions are shown.

### Tractography comparisons

As previously mentioned, if holes are present in tracking masks, tractography algorithms cannot track in this region. With *TractoFlow*, the lesions create GM voxels in the middle of the deep WM, which makes the tracking algorithm i) allowed to stop because there are falsely labeled GM voxels, and ii) prevents the tracking algorithm to explore through that region of WM. For example, Figure 5 shows a part of the corpus callosum segmented from the *TractoFlow* and *TractoFlow-ABS* tractogram. The extracted corpus callosum from *TractoFlow* contains broken streamlines by the lesions and lacks a large number of commissural streamlines, reconstructed by *TractoFlow-ABS*, going to the lateral left and right cortices.

## 4. Conclusion

*TractoFlow-ABS* uses the Freesurfer atlas to compute its tracking mask. As a result, in datasets with WM lesions, it is able to reconstruct a better WM mask, which allows tractography to track through the WM lesions if coherent local orientations exist. This improvement is thus mostly due to better tissue maps that come from the Freesurfer atlas. It is important to say that if one prefers the Freesurfer white matter segmentation than the FSL fast segmentation, it is possible to run this pipeline on healhty brains as well. *TractoFlow-ABS* is available on Github under license (https://github.com/scilus/TractoFlow-ABS).

